## Supplemental Figures for "Sequence Dependent Nanoscale Structure of CENP-A Nucleosomes"

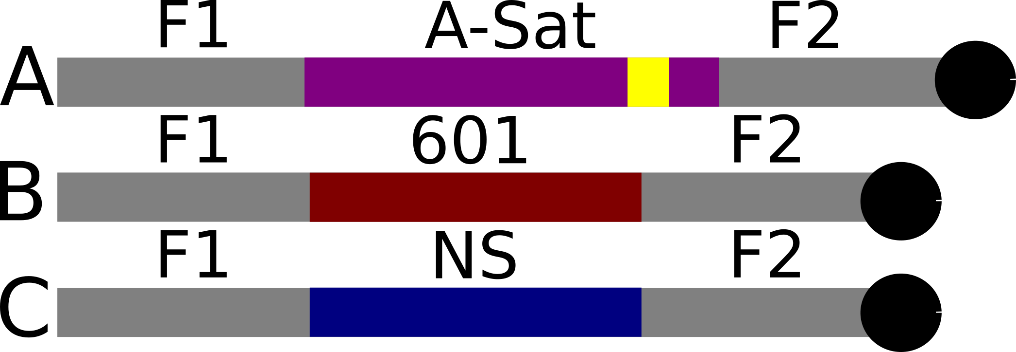

**Fig. S1.** Schematic representation of the substrates used. Schematic (A) represents the α-satellite substrate. Schematic (B) represents the 601 substrate. Schematic (C) represents the non-specific substrate. F1 is 114 bp and F2 is 125 bp on substrate (A). F1 is 113 bp and F2 is 117 bp on substrates (B) and (C). A-sat represents the α-satellite sequence of 171 bp. The yellow region represents the 17 bp CENP-B box within the α-satellite sequence. 601 represents the Widom 601 sequence of 147 bp. NS represents a non-specific sequence of 147 bp. The black circles represent the biotin label at the end of each template.

**Alpha-Satellite Sequence:**

5’GATGTGCTGCAAGGCGATTAAGTTGGGTAACGCCAGGGTTTTCCCAGTCACGACGTTGTAAAACGACGGCCAGTGAATTCGAGCTCGGTACCTCGCGAATGCATCTAGATGACCATTGGATTGAACTAACAGAGCTGAACACTCCTTTAGATGGAGCAGATTCCAAACACACTTTCTGTAGAATCTGCAAGTGGATATTTGGACTTCTCTGAGGATTTCGTTGGAAACGGGATAAAATTCCCAGAACTACACGGAAGCATTCTCAGAAACTTCTTTGTGATGAAGGGCGAATTCGAATCGGATCCCGGGCCCGTCGACTGCAGAGGCCTGCATGCAAGCTTGGCGTAATCATGGTCATAGCTGTTTCCTGTGTGAAATTGTTATCCGCTCACAATTCCACACAACATACG3’

**601 Sequence:**

5’GATGTGCTGCAAGGCGATTAAGTTGGGTAACGCCAGGGTTTTCCCAGTCACGACGTTGTAAAACGACGGCCAGTGAATTCGAGCTCGGTACCTCGCGAATGCATCTAGATGACACAGGATGTATATATCTGACACGTGCCTGGAGACTAGGGAGTAATCCCCTTGGCGGTTAAAACGCGGGGGACAGCGCGTACGTGCGTTTAAGCGGTGCTAGAGCTGTCTACGACCAATTGAGCGGCCTCGGCACCGGGATTCTCCAGGTCATCGGATCCCGGGCCCGTCGACTGCAGAGGCCTGCATGCAAGCTTGGCGTAATCATGGTCATAGCTGTTTCCTGTGTGAAATTGTTATCCGCTCACAATTCCACACAACATACG3’

**Non-Specific Sequence:**

5’GATGTGCTGCAAGGCGATTAAGTTGGGTAACGCCAGGGTTTTCCCAGTCACGACGTTGTAAAACGACGGCCAGTGAATTCGAGCTCGGTACCTCGCGAATGCATCTAGATGACCCTGGGGTGCCTAATGAGTGAGCTAACTCACATTAATTGCGTTGCGCTCACTGCCCGCTTTCCAGTCGGGAAACCTGTCGTGCCAGCTGCATTAATGAATCGGCCAACGCGCGGGGAGAGGCGGTTTGCGTATTGGGCGCTCTTCCGGACATCGGATCCCGGGCCCGTCGACTGCAGAGGCCTGCATGCAAGCTTGGCGTAATCATGGTCATAGCTGTTTCCTGTGTGAAATTGTTATCCGCTCACAATTCCACACAACATACG3’

**Fig. S2.** Sequences used for DNA substrates.

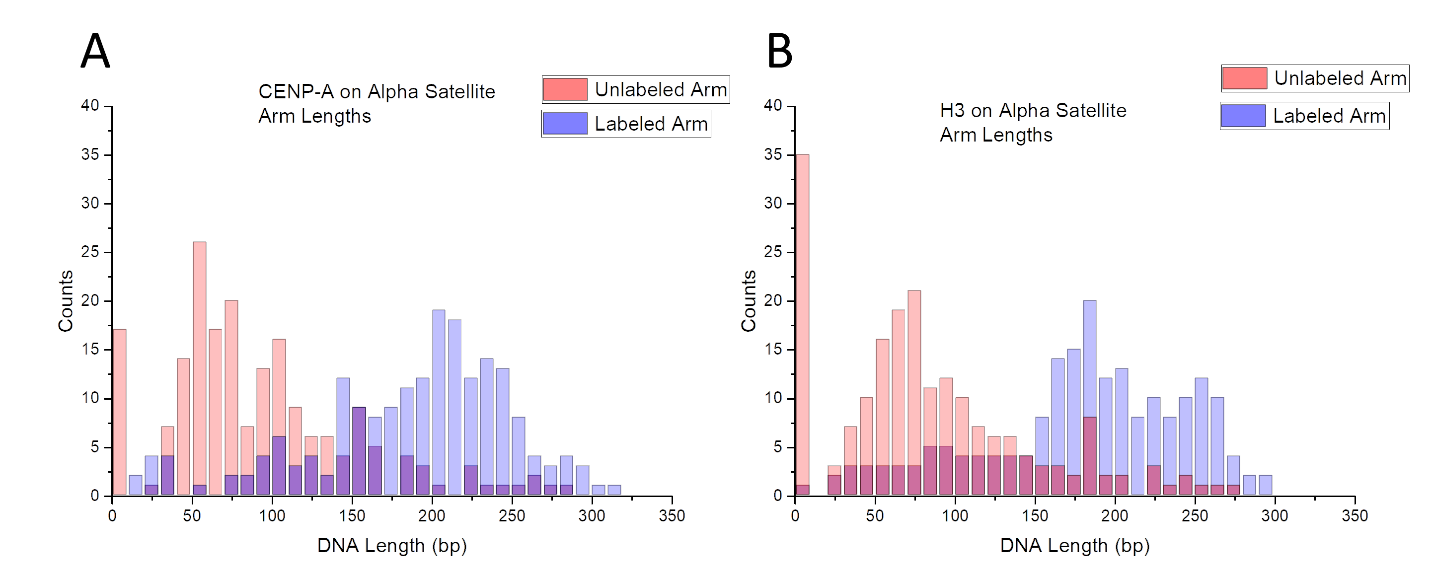
**Fig. S3.** Free DNA flank length measurements for nucleosomes assembled on the α-satellite substrate.

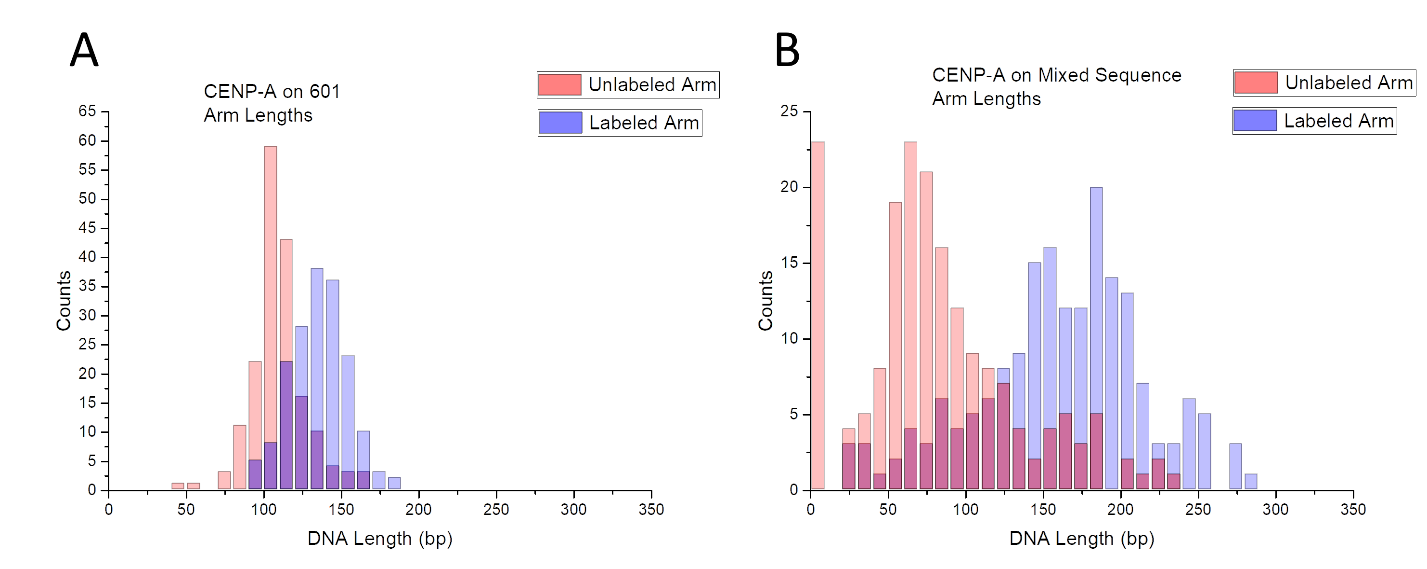

**Fig. S4.** Free DNA flank measurements for CENP-A nucleosomes assembled on the 601 motif **(A)** and the non-specific DNA **(B)**.
